# Landscape epidemiology of ash dieback

**DOI:** 10.1101/582080

**Authors:** M. Grosdidier, T. Scordia, R. loos, B. Marçais

## Abstract

Ash dieback caused by *Hymenoscyphus fraxineus*, an invasive alien pathogen, has been severely damaging European ash populations. Nevertheless, a large range of disease severities was observed at the landscape scale in the field. Several studies suggested that environment, such as climate, site conditions or local tree cover strongly affects ash dieback. We characterized the landscape epidemiology of the disease at two stages of the invasion process with spatio-temporal models using Bayesian models fitted by Integrated Nested Laplace Approximation (INLA). We first analyzed the effect of landscape features on the disease arrival and establishment stage at the scale of a village in NE France in 2012, 2 years after first report of the disease in the area and then, on the disease development stage in 2016-18. Landscape features had little impact on the disease at the establishment stage but strongly determined it further development. The local fragmentation of the tree cover was the most important factor with trees isolated or in hedges in agricultural settings far less affected then trees in forest environment. We showed that they were subjected to different microclimate with higher crown temperatures unfavorable to the pathogen development. Furthermore, host density was important for disease development with ash at low density far less affected by ash dieback. Presence of ashes in the vicinity affected local disease severity up to several hundred meters. These results may help to develop management strategy for the disease.

## Introduction

Ecologists have studied the interaction between spatial pattern and ecological processes in a landscape context since the 1960. However this research tread has been little explored in plant pathology, although the interest in landscape epidemiology developed in the last decade (*Plantegenest, Le May and Fabre* 2007; *Meentemeyer, Haas and Václavík* 2012). It has indeed been increasingly recognized that landscape features are important for pathogens spread, establishment and persistence in spatially structured host populations (*Hutchinson and Vankat* 1998; *Krewenka et al*. 2011; Kelly and *Meentemeyer*, 2002). Landscape epidemiology thus aims at determining how these mechanisms shape disease severity (*Plantegenest, Le May and Fabre* 2007). The spatial arrangement and composition of host vegetation is especially important for pathogens affecting long lived hosts such as forest trees (Condeso et al., 2007). For example, Perkins and Matlack (2002) suggest that the emergence of fusiform rust on Pines of south-eastern USA might have been promoted by a human induced change of the landscape with larger patches of host more favourable to the disease development. *Rodewald and Arcese* (2016) stressed out that landscape is especially important to explain the spread of invasive species, as some of it features can promote or hamper it dispersal. Roads or water courses can act as corridors for invasive species such as *Phytophthora* spp. because the propagules are dispersed by infected water or soil (*Jung and Blaschke* 2004; *Jules et al*. 2002). Infected fallen leaves can also be dispersed by vehicles as shown by the horse chestnut leafminer *Cameraria ohridella* (*Guichard and Augustin* 2002). Microclimate conditions that affected pathogen reproduction and survival are often determined by landscape pattern (Condeso et al., 2007; Kelly and *Meentemeyer*, 2002). Thus, topographic, edaphic, climatic, vegetation and historical landscape features can play a key role in disease expression and are often critical in the spread of invasive pathogens.

The relationship of disease with the landscape may be studied at several spatial and temporal scales using either static or dynamic models with the aim to respond to different questions such as inference or prediction. In order to do that, specific tools have been developed that enable to readily analyze data with spatial and temporal dependence. An increasingly used method, Bayesian space-time modelling approaches with Integrated Nested Laplace Approximation (INLA) is now widely available in R package. This tool has been used to analyze the effect of host structure or environment on disease presence or severity (*Schrödle and Held* 2011; *Marçais et al*. 2016).

Ash dieback is an emerging disease induced by an invasive pathogen, *Hymenoscyphus fraxineus* and has been severely damaging European ashes stands in the last decades. The fungus induces foliar infections, shoots blight and collar cankers that often lead to tree death. Originating from Asia, it was first reported in Poland in the nineties and spread across Europe to reach France in 2008. The biological cycle of *H. fraxineus* takes one year (*Gross et al*. 2014). Ash leaves are infected during the summer by airborne ascospores. At the end of autumn, infected leaves are dropped in the litter where the fungus overwinter in the leaf rachis under the protection of a black coloured structure called pseudosclerotial plate. In the spring, apothecia develop on the infected rachis and released ascospores in June-August. Asexual conidia, are involved in this sexual reproduction as they act as spermatia (*Gross et al*. 2012). *H. fraxineus* may remain and produce apothecia for 3-5 years on infected rachises in the litter (*Kirisits* 2015). At the end of summer, the pathogen extends from the leaves to the stem, inducing shoot mortality and significant crown dieback. This disease has been reported to threatened European ash populations and it was stressed that understanding how landscape might influence it development is of critical importance to well assess it impact and devise management strategies (Pautasso *et al*, 2013).

Several environmental factors such as soil moisture, air humidity, the proximity to water course, temperatures, stand age and stocking density that reflects canopy closure are known to influence disease severity (*Skovsgaard et al*. 2017). Reports on the impact of host density on the disease severity have been conflicting (*Skovsgaard et al*. 2017). Nevertheless, it is clear that dense stands with a closed canopy are very susceptible to the fungus while ashes set up in open habitats or near forest edges appears to be less affected (*Havrdová and Cerný* 2013; *Rosenvald et al*. 2015; *Vacek et al*. 2015). *H. fraxineus* has also been shown to have a poor survival at temperatures above 35°C, which affects the disease epidemiology (Hauptman *et al*. 2013, Grosdidier *et al*., 2018). The fungus also needs high moisture to develop, with moist sites deteriorating faster and showing more frequent canker at the trunk base (*Husson et al*. 2012; *Vacek et al*. 2015; *Marçais et al*. 2016; *Havrdová et al*. 2017).

Landscape feature could impact disease severity throughout multiple mechanisms, by promoting it spread (dispersal of infected rachises by road or river), affecting micro-climatic conditions important for it development or survival or through the density of ash host in the neighbourhood. To test these hypotheses, a landscape survey was realized at the scale of a village at different step of the disease development, either shortly after disease arrival or after several years of disease development. The effects of environment on crown dieback severity, collar canker and abundance of infected rachises in the litter were analysed in the frame of Bayesian hierarchical modelling.

## Material and Methods

### Sampling area

A study area of approximately 3.5 x 6.5 km was established around the village of Champenoux, in north east of France, at 15 km of Nancy (Lorraine) to encompass both forest and agricultural settings with scattered hedges and small woods in equal proportion. Ash dieback was observed in the area for the first time in 2010. Plots with a radius of 25 m were established on each node of a grid involving 737 nodes. The grid was of 200 x 200 m in forested area and of 141 x 141 m in agricultural settings to avoid over representing forest locations in the dataset. A first survey was done on aerial photography to control for the presence of trees on the nodes. Ground survey was then realized in 2012 on nodes with tree presence to check for the presence of ashes and to characterize the node environment. The number and size of ashes present on a 25m radius plot was recorded as well as presence of ash dieback. At that stage, samples were collected for analysis on 1-3 symptomatic ash per studied plot. The samples were analysed by qPCR for presence of *H. fraxineus* using the method developed by Ioos *et al*. (2009b).

A plot was set out in all nodes with presence of at least 5 ashes, whatever the size, which altogether represented 127 of the 797 nodes (Fig. 1).

**Fig. 1.**
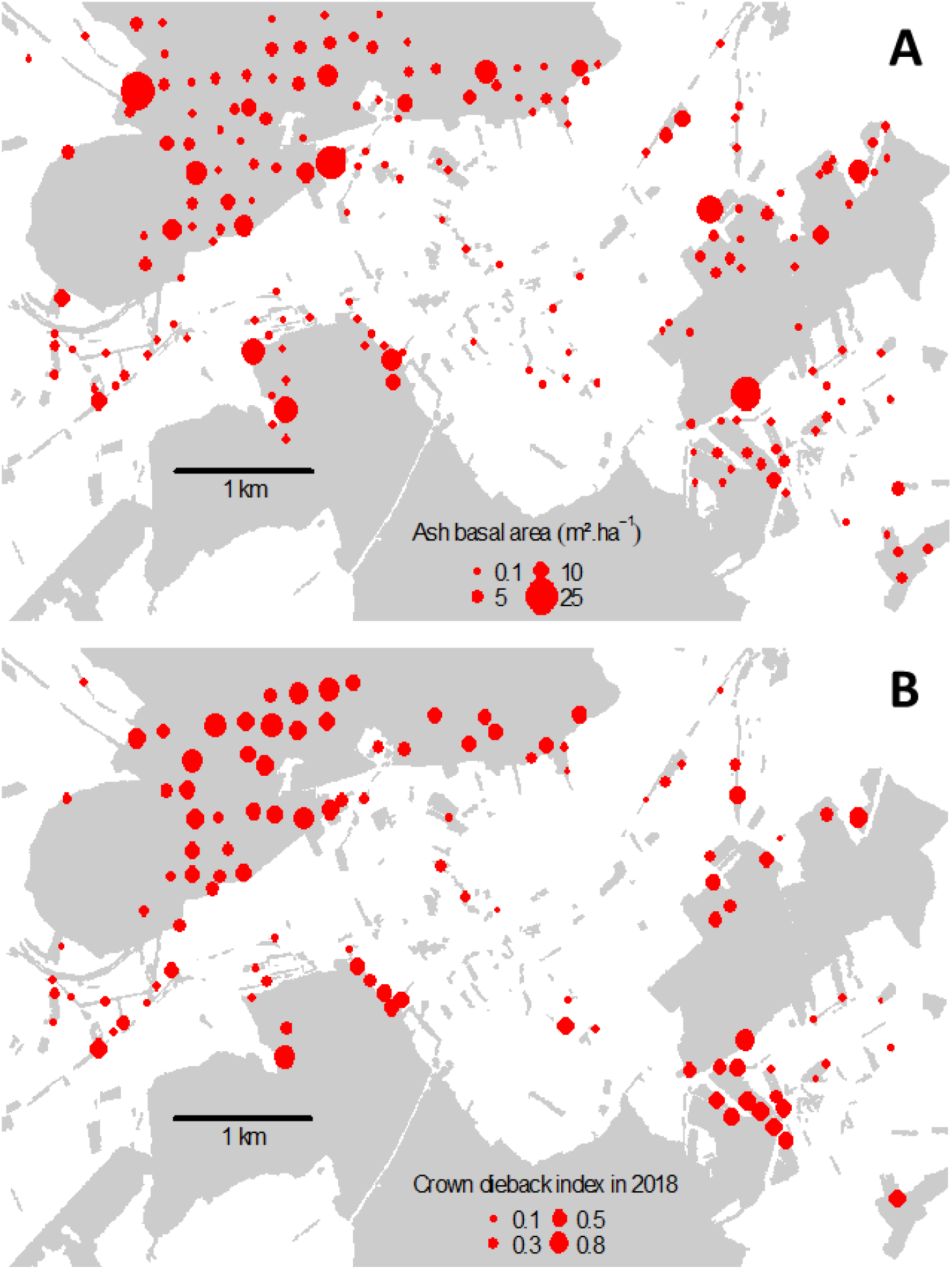
Plots spatial distribution in the studied area. (A) Host density, (B) Crown dieback index in 2018. The greyed area represents the tree cover, with large tracks of forested area in the north and south of the area.

### Environment characteristic of plots

Eighty plots were set up in forest while forty seven were in urban or agricultural environment. An index of tree cover fragmentation was computed from a tree cover shape file retrieved form the IGN site (http://professionnels.ign.fr). This shape file was corrected when needed from the aerial photographs showing the tree cover. The polygons of tree cover within a radius of 100 m from each plot were extracted and the perimeters to area ratio of those polygons were computed as an index measuring to which degree trees were isolated or within a tree stand. Distances of each plot from road (either coated road or dirt road), river and forest edge were computed with QGis software and Google earth. Computation was done in R software using the rgdal, rgeos and geosphere packages.

The basal area of ash present at each of the 127 plots was also characterized as a measure of host density. Three squares centred on the plot were set up with a diagonal of 16, 40 and 50 m oriented in cardinal directions (128, 800 and 1250 m^2^). The surface in which ash were sampled depended on their diameter at breast height (dbh). Ashes of dbh of 2.5 – 22.6 cm at were measured in the 128 m^2^ square, those of dbh of 22.6 – 37.6 cm in the 800 m^2^ square, and those of dbh over 37.6 cm in the square of 1250 m^2^. For each site, the ash basal area per ha was computed. To avoid over-estimating the basal area in areas with small woods, hedges or isolated trees, the total basal area was weighted by the proportion of tree cover within 100m from the plot. The ash basal area at node with too few trees to establish a plot (less than 5) was estimated using the initial count and dbh estimation of the ash trees present within the 25 m radius circle (Fig. 1a).

### Crown temperatures vs. forest environment

Temperate sensors EL-USB-2 | data loggers (Lascar Electronics Ltd UK, Wiltshire, United Kingdom) were set up in the crown of 16 ashes between the 9 July and the 12 September 2016. They were placed with a slingshot approximately in the middle of the crown and on the east side. Temperatures and relative humidity were measured each hour. Height ashes were isolates trees in agricultural settings while 8 ashes were in forest stands to compare the influences of the stand on the climatic conditions in the crown. The heights of the sensors, of the top and the bottom of the crown were determined. Hourly data of temperature were extracted of the Météo-France of Champenoux (WGS84 E 48.751 N 6.340) in 2016. Crown temperatures measured by data logger were modelled by a linear model depending on temperatures measured by Météo-France meteorological station of Champenoux at the same time and of the tree position (in forest stand or isolated tree). Hours were added as a factor in the linear model to take into account hourly variability in each day measured.

### Disease measurement

The first assessment of disease severity was realized in June - July 2012 (i.e. 3 years after first disease reports in the area) corresponding to the installation step of the disease. Three additional disease assessments were realized in June 2016, June 2017 and June 2018 (i.e. 7-9 years after first disease reports) which corresponds to development step of the disease. Ashes crowns were rated according to 6 dieback scores: 0 for no dieback, 0.05 for 1 to 10 % of dieback, 0.35 for 10 to 50% of dieback, 0.625 for 50 to 75% of dieback and 0.875 for 75 to 99% of dieback or tree with stem dead and sprout at the stem base, 1 for trees dead. Crown dieback index of each plot was computed by averaging the dieback score of all ash of the plot. Moreover, the presence or absence of a canker at the trunk base was observed for each of the studied tree. While plots located on the border nodes were characterized for the environment and the host density, they were not observed for disease intensity. Depending on the year, 100 to 114 plots were rated, some plots being abandoned because no ashes were present anymore while others were added because of ash recruitment.

To better characterize the inoculum sources, ash rachis density in the litter was determined for each of the studied plots in May 2016 and from Mars to May 2017. All rachis on a 10 cm wide band along a 10 m transects were sampled in each plot. In 2016, one 10 m transect was sampled. In 2017, two 10-m perpendicular transects were sampled. The rachises were sorted in the laboratory in 2 categories: either infected i.e. black rachis with presence of a pseudosclerotial plate, or healthy, i.e. white or beige rachis without any pseudosclerotial plate. In 2016, we determined the length of rachis in each of those 2 categories. In 2017 the dry weight in each of the 2 categories was determined after 48 hours in a heat chamber at 50°C. The density of rachises in each category was computed in m.m^−2^ of litter using a linear equation between the 2 measurements established in 2016 (L = 163.73*DW, with L, the length in cm and DW, the dry weights of rachises (in *Grosdidier*, unpublished results). The validity of the rachis infection assessment was checked in 2016. Infected and non-infected rachis were sampled (20 of each per plot on 28 plots) and placed in laboratory conditions in a wet chamber at 20°C during about 1 month to assess their ability to produce *H. fraxineus* apothecia

Additionally, infected rachises were harvested in the litter in August along 3 areas of 1 x 0.1 m each in 23 plots in 2016 and 31 in 2017. Eleven plots were visited the both years. Length of infected rachises was measured in the laboratory as previously described and the number of apothecia present on each of them was counted. Number of apothecia per length of infected rachises was then determined. The sampled plots were selected to cover the available range of tree cover fragmentation and distance to the river. The log transformed nb of apothecia per cm of infected rachis was analysed by regression using the year, the tree cover fragmentation and the log of distance to the river as independent variables.

### Statistical analysis

Statistical analyses were done separately for the installation stage of the disease in the area corresponding to 2012 data, and for the disease development stage corresponding to 2016-18 data.

The crown dieback index, proportion of infected rachises and proportion of collar canker were analysed using R package “INLA” to account for the spatial and temporal dependencies. *y_it_* was either the mean proportion of crown dieback, the proportion of tree with collar canker, the proportion or the total density of infected rachises in the litter at plot *s_i_*(*i* = 1,…,*n*) and in year *t* = 1,…,*T*. To analyse the crown dieback index which corresponds to proportions not based on number of cases observed, a 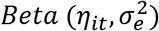 distribution was used. For the collar canker prevalence or the proportion of infected rachises, a 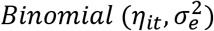 distribution was assumed. Lastly, the density of infected rachises in the litter was log transformed and analysed with a 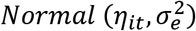 distribution. 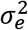 represents the variance of the measurement error defined by a Gaussian white noise process, both serially and spatially uncorrelated, and 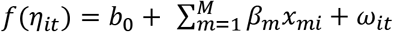, where *b*_0_ is the intercept, and *β*_1_,…,*β_M_* are the fixed effects related to covariates *x*_1_,…,*x_M_*. The link function f used was the logit function for the mean proportion of crown dieback, the proportion of tree with collar canker, the proportion of infected rachises in the litter and the identity function for the total density of infected rachises in the litter. The term *ω_it_* refers to the latent spatio-temporal process which changes in time with a first-order autoregressive dynamics and a spatially correlated innovations: *ω_it_* = *aω*_*i*(*t*−1)_ + *ξ_it_*, with *t* = 2,…,*T*, |*a*| < 1 and 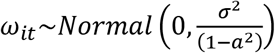. Moreover, *ξ_it_* is a zero-mean Gaussian field, assumed to be temporally independent and characterized by the following spatio-temporal covariance function: 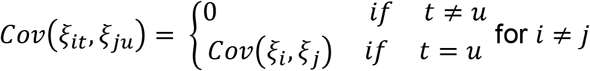, where *Cov*(*ξ_i_,ξ_j_*) is given by a Matern spatial covariance function defined by 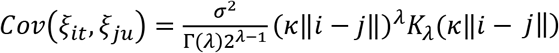, where ||*i*−*j*|| is the Euclidean distance between two generic locations and *σ*^2^ is the marginal variance. *K_λ_* denotes the modified Bessel function of the second kind and order *λ* > 0, which measures the degree of smoothness of the process and is usually kept fixed due to poor identifiability. Conversely, *κ* > 0 is a scaling parameter related to the range 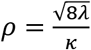 which corresponds to the distance at which the spatial correlation is close to 0.1, for each 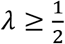. A mesh for the spatial domain was first created.

Analysis of at the disease installation stage was realized with no temporal variation in the spatial field. Thus, term *ω_it_* corresponds only to *ξ_it_* with t = 2012. The full spatio-temporal model was compared to simplified versions of the model: (i) a spatial model only with no first-order autoregressive dependency between years (*ω_it_* equals to *ξ_it_* for t = 1 for 2016, 2 for 2017 and 3 for 2018); (ii) a temporal model only with no spatial dependency; (iii) a spatio-temporal model with independent spatial and temporal effect. Models were compared using the Deviance Information Criterion (DIC). The model with the lowest DIC was chosen to assess for effect of covariates.

In a preliminary step, the range at which host density might impact ash dieback was studied. The hypothesis was that host patches might enhance the disease development depending on their host basal area and on the distance to the focal point. Two kernels were tested to determine how this influence may decay with distance, the exponential and the Epanechnikok kernel, using ranges of 0,100,200,300,400 and 500. The host abundance in the neighbourhood of a plot i (HAN) was then computed as 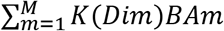 with BA, the ash basal area at plot m, dim, the distance between plot I and m and K, the kernel. To account for a density of the grid that was twice in agricultural settings, BA_m_, the ash basal area in neighbouring plot m located in those agricultural settings was divided by 2. Full spatial models were then fitted using these computed host abundance in the neighbourhood with 2018 data only and the best kernel and appropriate range were selected according to the DIC. This variable HAN was then used as a measure of local host density.

Full spatial / spatio-temporal models were built with a forward strategy for entering variables in the modelCovariates selected for entry was the one with fitted parameter with 2.5% and 97.5% quantile ranges that did not span zero and yielding the lower DIC.

## Results

### Sites characteristics and *H. fraxineus* presence

Altogether, depending on mortality, logging and recruitment of ashes during the study period, 100 plots were observed in 2012, 100 in 2016, 109 in 2017 and 111 in 2018. Most of the plots were located in forest environment, with 67 plots in forest settings and 44 in agricultural ones in 2018. However, many plots had intermediates conditions being either located at forest edges or in small woods within agricultural settings (15 forest plots and 16 plots in agricultural settings in 2018). To account for this, we used the tree cover fragmentation index that was indeed higher in agricultural settings (0.028 ± 0.003) than in forests settings (0.153 ± 0.019). Ash basal area was significantly higher in forest than in agricultural settings (p-value < 0.01, Fig.1a), with mean values of ash basal area of 6.1 ± 1.2 m^2^.ha^−1^ in forest and 3.3 ± 1.2 m^2^.ha^−1^ in agricultural settings. Only 2 pure ash stands were present in forest environments of the area. Ash was usually at low density in stands mixed with oaks and hornbeam. By contrast, small woods in agricultural settings although limited in size were often pure ash stands.

In 2012, only 2 years after disease arrival in the village of Champenoux, ash dieback was observed in all studied plots with 74% of the ashes showing at least limited disease symptoms and 47% of them showing moderate to severe crown decline. The mean crown dieback index was 28 ± 3%. *H. fraxineus* could be identified by qPCR from 107 of the 115 sampled plots (93%). The plots where *H. fraxineus* could not be detected by qPCR were stands with 1-3 larges ashes trees where adequate samples were difficult to collect. Nevertheless, only six trees among the 1424 trees observed in 2012 showed presence of a collar canker. In 2016, 4 years after, the disease had increased in severity (Fig. 2a, b). Mean crown dieback index observed in 2016, 2017 and 2018 were of 36 ± 3, 33 ± 3 and 40 ± 4% respectively. Collar cankers were observed on respectively 231, 229 and 294 of the trees in 2016, 2017 and 2018 (16%, 18.4% and 25.1% respectively).

**Fig. 2.**
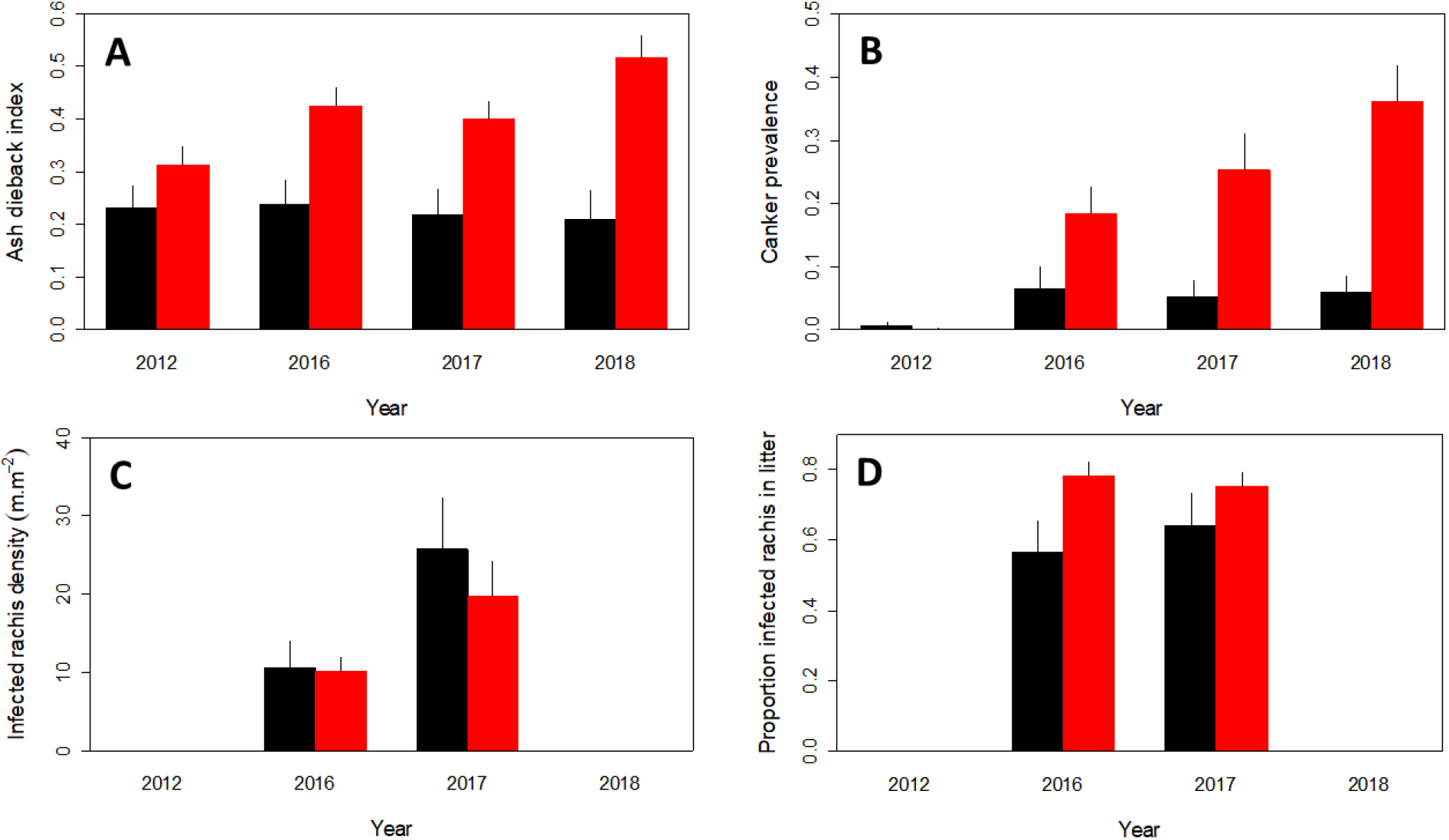
Evolution of ash dieback in the studied plots from 2012 to 2018. A. Ash dieback index (proportion of shoot mortality), B. Collar canker prevalence, C. Density of ash rachis infected by *H. fraxineus* in the litter, D. Proportion of ash rachis in the litter that are infected by *H. fraxineus*. Black bar: agricultural settings: Red bar, forest settings.

### Effect of the landscape on ash dieback, collar canker and rachis infection

In a preliminary step, the range at which host density might impact crown dieback was studied. The host abundance in the neighbourhood (HAN) computed using an exponential kernel with a range of 200 m yielded the lower DIC for both the 2018 ash dieback index and basal canker prevalence (Fig. 3). The respective DIC were of −128.7 and 355.9 compared to −116.5 and 363.2 for the plot ash basal area. Using the Epanechnikof kernel for computing the HAN yielded slightly larger DIC.

**Fig. 3.**
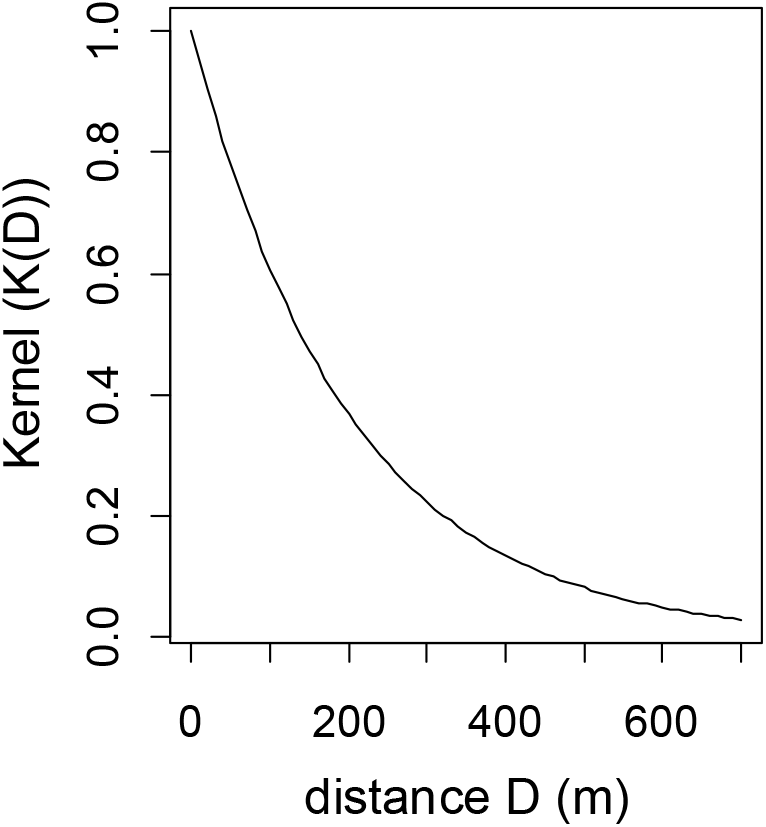
Decay with distance of influence of the host abundance in the neighbourhood on ash dieback. The best kernel was the exponential kernel with a range of 200 (K(G) = exp(-D / 200)).

#### Disease establishment step (2012)

The analysis at the installation step of the disease was done only for the crown dieback index, collar cankers being very infrequent. The only variables that showed a significant relationship with crown dieback index in 2012 were the tree size (dbh) and the tree cover fragmentation (Table 1). The dieback index decreased with tree dbh, smaller ashes being more severely affected, and also with the tree cover fragmentation. Ashes in closed canopies were more severely affected than ashes in hedges or isolated ashes. Hence, since 2002, crown decline was less severe in agricultural settings where tree cover is more fragmented compared to forest (Fig. 2a). At that stage, host abundance (HAN) and proximity of the plot to river or road were not linked to the dieback severity (Table 1). The estimated range of the spatial dependency in crown decline was 542 m. The spatial field explained some variability in the data that was not taken into account by covariates (Fig. 5).

**Table 1:** Effects of landscape features on ash crown dieback at the arrival and development stage of the disease. Significant values are in bold.

| Variable<sup>a</sup> | Establishment stage (2012) |  | Development stage (2016 – 18) |  |
| --- | --- | --- | --- | --- |
|  | Mean (sd) | P2.5 ; P97.5 | Mean (sd) | P2.5 ; P97.5 |
| Host abundance in neighbourhood (HAN) | 0.01 (0.01) | -0.01 ; 0.03 | **0.04 (0.01)** | 0.02 ; 0.06 |
| Tree cover fragmentation | **-2.64 (1.14)** | -4.96 ; -0.46 | **-7.04 (1.12)** | -9.24 ; -4.84 |
| Trees size (dbh) | **-0.02 (0.01)** | -0.04 ; -0.01 | **-0.02 (0.004)** | -0.03 ; -0.01 |
| log(Distance to river) | -0.03 (0.05) | -0.13 ; 0.08 | **-0.13 (0.04)** | -0.20 ; -0.05 |
| log(Distance to road) | -0.04 (0.05) | -0.13 ; 0.05 | 0.04 (0.03) | -0.10 ; 0.04 |
| $\sigma_e^2$ | 0.116 (0.018) | 0.009 ; 0.158 | 0.057 (0.007) | 0.045 ; 0.071 |
| $\kappa$ | 0.010 (0.011) | 0.002 ; 0.004 | 0.005 (0.002) | 0.002 ; 0.009 |
| $\sigma^2$ | 0.030 (0.040) | 0.001 ; 0.143 | 0.516 (0.158) | 0.276 ; 0.892 |
| $\rho$ | 542 (356) | 69 ; 1374 | 645 (228) | 302 ; 1189 |
| DIC Null model | -103.6 |  | -466.1 |  |
| Full model | -110.9 |  | -508.1 |  |

<sup>a</sup> Host abundance in neighbourhood, sum of Ash basal area in neighbouring plots weighted by the exponential kernel with a parameter of 200, Tree cover fragmentation, Perimeter / area ratio of the tree cover within 100 m.

#### Disease development step (2016-18)

Models without covariates with only a spatial effect, a temporal effect, both a spatial and a temporal effect without time * space interactions were compared to the full spatio-temporal model. For crown dieback index, the respective DIC were of −445.7, −151.7, −463.5 and −466.0 indicating that the model with time, spatial effect and their interactions had a better fit, although the time * space interactions brought just a small improvement. The same results were obtained with canker prevalence and proportion of rachis infected (result not shown). Spatio-temporal models with interaction between spatial and temporal effects were thus chosen for further analysis.

The crown decline was more strongly influenced by landscape at the development step. Tree cover fragmentation, ash abundance in the neighbourhood (HAN) and distance to river all were significantly related to the both the crown dieback index and collar canker prevalence (Table 1 and 2). As during the installation step, ash located in plot with a higher tree cover fragmentation index were far less affected by ash dieback (Fig. 1b, Fig. 4a). Strikingly, the diseased overall increased in severity with time in forest conditions but not in agricultural settings, where tree cover was highly fragmented (Fig. 2 a, b). Ash dieback strongly increased with HAN (Fig. 4b). Plots with low a HAN remained relatively healthy. Ash dieback severity also decreased with distance to river (Table 1 and 2). Additionally, the crown dieback index significantly increased with tree size (mean plot dbh) and with distance to the road (Table 2, Fig. 4). The range of the spatial correlation was in a similar range than at the disease arrival step and corresponds to 645 m for crown decline and 403 m. The spatial effect showed similar patterns across the years (Fig. 5).

**Fig. 4.**
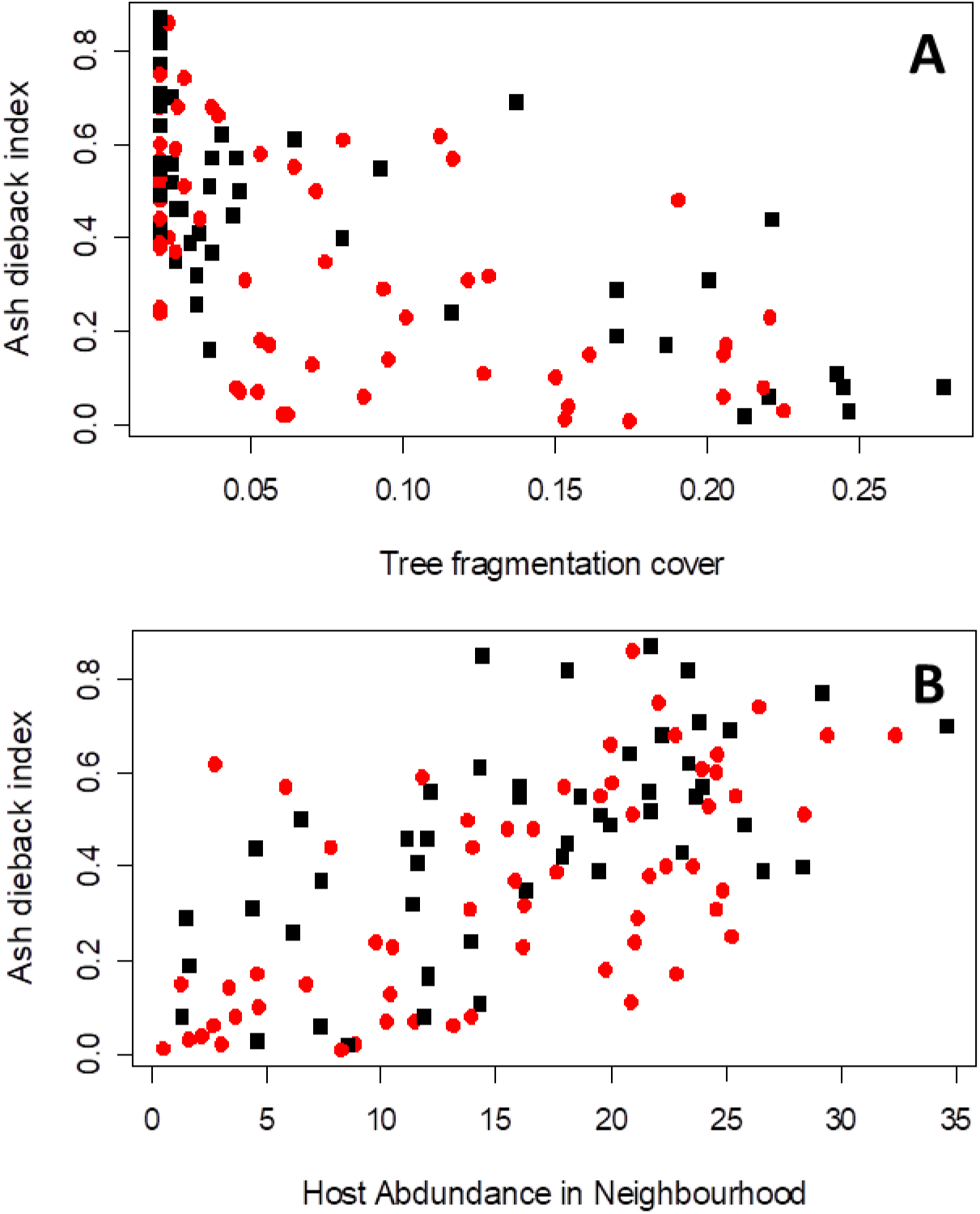
Factor controlling the severity of ash dieback. Effect of (A) tree cover fragmentation and (B) host abundance in the neighbourhood (HAN) on the crown dieback index. The black squares represent sites with mean tree trunk diameter less than 15 cm while the red circles are those with mean tree trunk diameter over 15 cm.

**Fig. 5.**
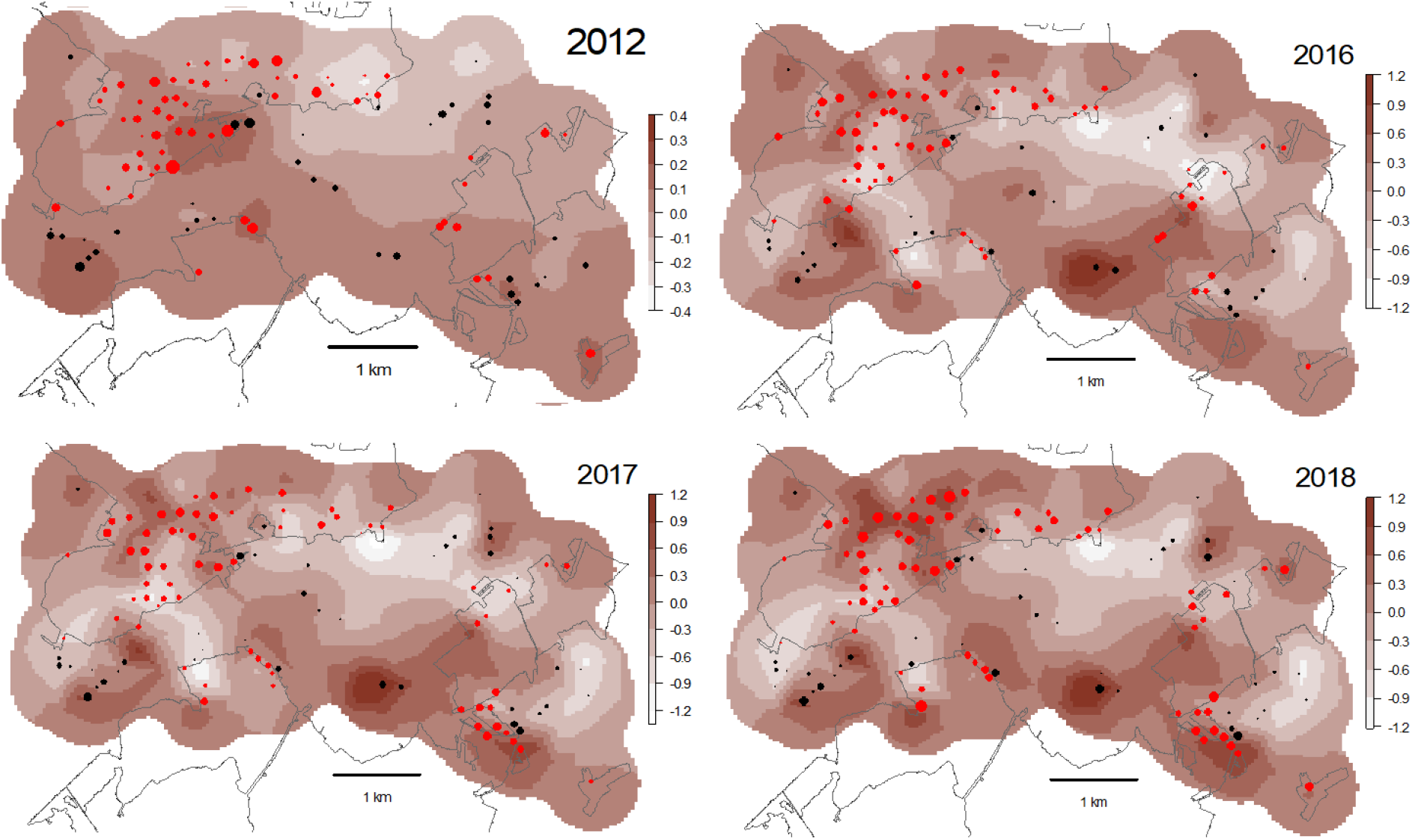
Map of the posterior mean of the spatial random fields for ash dieback model for the disease establishment (2012) and development steps (2016-18). Red dots are forest settings while black dots are agricultural settings. Dots size corresponds to ash dieback severity.

**Table 2:** Effects of landscape features on prevalence of collar canker and % infected rachises in the litter at the development stage of the disease (2016 – 2018). Significant values are in bold.

| Variable<sup>a</sup> | Prevalence of collar canker |  | % infected rachises in litter |  |
| --- | --- | --- | --- | --- |
|  | Mean (sd) | P2.5 ; P97.5 | Mean (sd) | P2.5 ; P97.5 |
| Host abundance in neighbourhood (HAN) | **0.06 (0.02)** | 0.03 ; 0.10 | **0.05 (0.01)** | 0.02 ; 0.08 |
| Tree cover fragmentation | **-7.53 (2.46)** | -12.36 ; -2.71 | -0.92 (2.12) | -5.09 ; 3.25 |
| Trees size (dbh) | 0.01 (0.01) | -0.01 ; 0.03 | -0.012 (0.008) | -0.03 ; 0.01 |
| log(Distance to river) | -0.002 (0.08) | -0.17 ; 0.16 | **-0.23 (0.06)** | -0.51 ; -0.11 |
| log(Distance to road) | **0.135 (0.07)** | 0.01 ; 0.27 | 0.03 (0.06) | -0.09 ; 0.14 |
| $\sigma_e^2$ | 0.327 (0.048) | 0.257 ; 0.445 | 0.366 (0.057) | 0.286 ; 0.509 |
| $\kappa$ | 0.008 (0.002) | 0.004 ; 0.014 | 0.009 (0.003) | 0.005 ; 0.015 |
| $\sigma^2$ | 3.027 (0.933) | 1.678 ; 5.313 | 4.382 (1.241) | 2.575 ; 7.411 |
| $\rho$ | 403 (117) | 208 ; 664 | 352 (97) | 185 ; 560 |
| DIC Null model | 1050.0 |  | 987.0 |  |
| Full model | 1029.4 |  | 973.1 |  |

<sup>a</sup> Host abundance in neighbourhood, sum of Ash basal area in neighbouring plots weighted by the exponential kernel with a parameter of 200, Tree cover fragmentation, Perimeter / area ratio of the tree cover within 100 m.

Fructification tests shown that *H. fraxineus* produced apothecia on very few of the rachises classified as non-infected (0.9%). Usually just a very short section of the rachis was then observed to produce apothecia. By contrast, *H. fraxineus* produced apothecia on the large majority of the rachis classified as infected (97%). Thus, the visual assessment of rachis infection was valid. Global proportions of infected rachises observed in the litter were very high, at 70-75% in both 2016 and 2017 (range 20-100%), even in locations with little crown dieback (Fig. 2d). The proportion of infected rachises in the litter increased with host density (HAN) and proximity to the river (Table 2). By contrast, it was not significantly influenced by tree cover fragmentation, although it was marginally higher in forest settings than in agricultural settings (Fig. 2d). The range of spatial correlation was in a similar order of magnitude compared to crown decline and collar canker prevalence at 352 m. Total length of infected rachis in the litter is of special interest as it is a measure of the potential for inoculum production. The density of infected rachis par plot was highly variable, with median values of 8 and 14 m.m^−2^ in 2016 and 2017 (range of 0.2-60 m.m^−2^, Fig. 2c). It increased with host local density (plot ash basal area, 0.03, confidence interval CI [0.01, 0.06]) and mean ash size (tree dbh, 0.02, CI [0. 1, 0.04]). It was however not significantly related to any of the studied landscape features, in particular to the tree cover fragmentation (parameter 2.2, CI [-0.5, 5.1]). Density of infected rachis in the litter was in similar range in forest and agricultural settings (Fig. 2c). The number of apothecia per m of infected rachises in early July was 68 ± 14 in 2016 and 47 ± 17 in 2017. Apothecia abundance increased significantly close to river and was higher in 2016 (both p-values < 0.01), but did not depend on tree cover fragmentation (p-value= 0.101). Hence, the amount of *H. fraxineus* apothecia per length of infected rachis was similar in forest and in agricultural settings.

### Temperature modelling

Temperatures data logger set up in the ash crown in summer 2016 showed that temperatures in the crown of isolated trees in agricultural settings were above those of forest trees by 0.4°C in average and were about more 4h longer above temperatures of 35°C (p-value < 0.05). Height of data logger in the crown did not influence the results (p-value = 0.34).

The model developed to explain hourly temperature in the tree crown accounted for 91% of variability observed (p-value < 0.01). In order to investigate the possible influence of the heat wave that occurred in Eastern France in 2015, averages of maximum daily temperatures inferred by the model in 2015 and 2016 were used for further analysis. We computed for each year the number of hour where crown temperature reaches values above 35°C. Modelled temperatures were seldom higher than 35°C in 2016 in both forest and agricultural settings (0 – 11h depending on the settings and the year). By contrast, in 2015 modelled crown temperature were above 35°C during 24h in forest and during 64h in agricultural settings.

## Discussion

Our results illustrate how landscape can affect the colonisation and subsequent development of an invasive disease in a new area. While initial colonisation was not strongly affected at the scale of the studied area, several landscape features, in particular host density and tree cover fragmentation, were very important for the latter development of the disease. Further, circumstantial evidence indicated that high summer temperature may significantly limit the disease development, even in a temperate climate as NE France.

The colonisation of the studied area was very quick, the pathogen being present everywhere 2 years after it first report. Disease severity could not be related to potential dispersal corridors such as tracks of dense host populations or road and river that might have dispersed infected rachis. The relation between environmental heterogeneity, in particular host distribution, and the ability to of a pathogen to invade an area have been well documented both theoretically and empirically (Park *et al*. 2001, Kaufmann and Jules. 2006, Condeso et al. 2007, Pautasso and Jeger, 2008). However, it would appear that the fragmentation of the host population did not limit the ability of *H. fraxineus* to colonise the area given it very efficient aerial dispersal (Gross *et al*. 2014). The mean range for *H. fraxineus* ascospores dispersal from inoculum sources has been shown to be as high as 1.5 to 2.5 km (Grosdidier *et al*. 2018). With such a dispersal range, the landscape of NE France and of many other regions of Europe may be perceived by the colonising *H. fraxineus* as a continuous host cover. All patches of Ash within the studied area were colonised since 2012 and remained so during the study period despite the occurrence of a heat wave in 2015 that was unfavourable to the pathogen expression. This is probably related to the ability of *H. fraxineus* to survive in rachises of the litter during several years which gives some stability to the pathogen population (Kirisits, 2015). Some landscape features could however be related to disease severity since the colonisation stage, in 2012 (tree cover fragmentation and tree size). As these two features remained important throughout the studied period, it can be argued that they influenced mostly local disease intensification. It has for example already been reported that small ashes are more affected by *H. fraxineus* (*Husson et al*. 2012; *Skovsgaard et al*. 2017). The pattern of spatial correlation (spatial random fields for ash dieback model) observed in 2012 shows similarities with what was observed in later years which might reflect patches of earlier colonisation by *H. fraxineus*. Alternatively it could represent underlying environment heterogeneity not captured by the measured variables or heterogeneity in the resistance level of ashes populations (*McKinney et al*. 2014).

Although not important for the colonisation stage, environmental heterogeneity proved to be critical for the disease subsequent development. In particular, we demonstrated the importance of host density for ash dieback. While considered critical in disease dynamic, this parameter has not been documented often in disease of plants in wild ecosystems (Plantegenest et al, 2007, Keesing *et al*. 2010). Experimental work has demonstrated it importance in both grassland and trees systems (Knops *et al*. 1999; Mitchell *et al*. 2002, Borer *et al*. 2009; Hantsch *et al*. 2013, Parker and Gilbert. 2018), but well documented empirical studies remain scarce (Haas *et al*. 2011). However, it has to be pointed out that host density dependant tree pathogens have been shown to promote diversity in tropical forest by maintaining individual species at low density, lowering the overall inter-species competition (so called Jenzen-Connell effect, Bever *et al*. 2015). We showed that presence of large ash populations in the neighbourhood has an influence on ash dieback that decayed exponentially but was still significant 200-300 m away. The index of host abundance used, HAN can be seen as a surrogate for the force of infection: the density of infected ash rachises in the litter was strongly related to the local host density and tree size, but not to the studied environmental parameters while apothecia production on the infected rachises increased only in water courses vicinity. Thus, inoculum production depended mainly on ash basal area. The range of the spatial dependency to neighbouring ash populations matches the range of ascospores dispersal (Grosdidier et al, 2018). Although *H. fraxineus* ascospores can be detected up to 500 m from inoculum sources, the spore load in the air drops quickly between 0 and 50 m. Analysis showed that incorporating HAN in the model did not account for all spatial dependency; some remained at a range of 300-600 m depending on the disease variable considered, which could reflect uncomplete account of the Ash density on the used grid design. The strength of the host density effect can be explained by the disease aetiology. Asexual secondary infection has been reported to be of limited epidemiological importance (Gross et al, 2012). As *H. fraxineus* is a not systemic pathogen, it probably requires massive leaf infection to produce a dieback. This may be even more relevant for collar cankers that appear to need large inoculum load to occur (Marçais *et al*. 2017). Indeed, the leaf infection appears to be massive: the proportion of infected rachises in the litter was usually very high (about 70%, with minimal values of 20%) and depended on host density and proximity to river. The host density effect we demonstrated for ash dieback could be used for devising management strategies. In the landscape of NE France, Ash is often at low host density, either because it is located in agricultural settings as hedges or isolated trees or because it is often present in forests as a minor species in mixt stands together with oaks or beeches. This study shows that promoting trees diversity in forest stand may be a valuable strategy to reduce their vulnerability to an invasive pathogen such as *H. fraxineus*. We are experiencing an increasing rate of harmful forest pest invasions (Santini *et al*. 2013).Moreover, they remain unpredictable because the pathogens are usually described late in the invasion process. Thus, acting on tree density to lower forest stands vulnerability to future invaders appears attractive because it is a non-targeted strategy.

Forest settings were much more favourable to ash dieback than agricultural settings. This could partly be because agricultural settings often have lower ash density. However, this cannot be the only explanation as both the length of infected rachis in the litter and the production of apothecia per length of infected rachis did not depend on tree cover fragmentation. Alternatively, it could be relate to micro-climatic conditions more favourable to *H. fraxineus* in forests. It is known that closed canopies create more humid and cool micro-climate (*Villegas et al*. 2010), both conditions known to be favourable to leaf pathogen. Indeed, crowns of isolated trees were more frequently exposed to temperature above 35°C which are very unfavourable for *H. fraxineus* survival (*Hauptman et al*. 2013). It has been shown that leaf temperature can be above air temperature by several degrees Celsius. High site moisture was also favourable to ash dieback, as both crow dieback and inoculum production increased in proximity to rivers. Several authors showed that high moisture promote ash dieback (*Husson et al*. 2012; *Marçais et al*. 2016) and it has been hypothesized that dense stocking could favour disease severity by enhancing air humidity (Havrdová *et al*. 2017). *Rosenvald et al*. (2015) showed trees in stand edge were less affected by *H. fraxineu*s and hypothesised that high temperatures could limit the disease development in these environments. Last, high summer temperature were demonstrated to limit ash dieback was in the Rhône valley in France at the southern limit of the disease presence (*Grosdidier et al*. 2018). We show here that crown temperatures may reach levels that limit ash dieback during heat wave in a temperate continental climate such as NE France. Interestingly, higher ash dieback severity in closed canopies was not related to higher inoculum production potential, suggesting that leaf infection was less strongly affected by tree cover fragmentation. However proportion of rachis infected in the litter could be only loosely related to leaf infection because infected rachis persist several years in the litter and may decompose at rate different than none infected rachis. The lower basal canker prevalence in fragmented canopies has significant practical interest. Basal cankers induced by *H. fraxineus* have been associated with secondary infection by root rotting fungus such as Armillaria which drastically decrease tree stability (Marçais *et al*. 2016, Skovsgaard *et al*. 2017). We found not only that those basal cankers are far less present in hedges or isolated ashes that are frequent on road sides, but that their prevalence decreases in road vicinity.

Ash dieback has been reported to threaten the European ash population as well as some species that depends exclusively on this tree species (Pautasso *et al*. 2013, Mitchell *et al*. 2014). This assessment, made early in the development of the pandemic in Europe, appears based on the extrapolation of worst observed situations to the entire European ash population. We show here that this might be overly pessimistic as environmental heterogeneity at the landscape level should strongly limit the global disease severity. A large number of ash populations are in situations far less exposed to ash dieback, being either in fragmented tree covers or at low host densities. Another positive point is the significant effect of high summer temperatures during heat waves which appears to have limited disease severity in the studied area. With the predicted global warming, the frequency of summer temperatures above 35°C might increase in the area, which might be favourable to ash populations, at least in the near future.

## Acknowledgements

We thank Louis Cordonnier, Yann Guépet, Aurélie Backes, Pauline Hehn, Amrane Chabane-Chaouche, Anaïs Gillet and Olivier Caël for their technical assistance. This work was supported by grants from the Forestry Health Department, French Ministry of Agriculture and Forestry, ANSES. The UMR1136 research unit is supported by a grant managed by the French National Research Agency (ANR) as part of the “*Investissements d’Avenir*” program (ANR-11-LABX-0002-01, Laboratory of Excellence ARBRE).

